# Brain Differences in the Prefrontal Cortex, Amygdala, and Hippocampus in Youth with Congenital Adrenal Hyperplasia

**DOI:** 10.1101/791541

**Authors:** Megan M. Herting, Anisa Azad, Robert Kim, Mitchell E. Geffner, Mimi S. Kim

## Abstract

**Context:** Classical Congenital Adrenal Hyperplasia (CAH) due to 21-hydroxylase deficiency results in hormone imbalances present both prenatally and postnatally that may impact the developing brain.

**Objective:** To characterize gray matter morphology in the prefrontal cortex and subregion volumes of the amygdala and hippocampus in youth with CAH, compared to age- and sex-matched controls.

**Design:** A cross-sectional study of 27 CAH youth (16 female; 12.6 ± 3.4 year) and 35 typically developing, age- and sex-matched healthy controls (20 female; 13.0 ± 2.8 year) with 3-T magnetic resonance imaging scans. Brain volumes of interest included bilateral prefrontal cortex, and eight amygdala and six hippocampal subregions. Between-subject effects of group (CAH vs control) and sex, and their interaction (group-by-sex) on brain volumes were studied, while controlling for intracranial volume (ICV) and group differences in body mass index and bone age.

**Results:** CAH youth had smaller ICV and increased cerebrospinal fluid volume compared to controls. In fully-adjusted models, CAH youth had smaller bilateral, superior and caudal middle frontal volumes, and smaller left lateral orbito-frontal volumes compared to controls. Medial temporal lobe analyses revealed the left hippocampus was smaller in fully-adjusted models. CAH youth also had significantly smaller lateral nucleus of the amygdala and hippocampal subiculum and CA1 subregions.

**Conclusions:** This study replicates previous findings of smaller medial temporal lobe volumes in CAH patients, and suggests that lateral nucleus of the amygdala, as well as subiculum and subfield CA1 of the hippocampus are the most affected regions in CAH youth.

**Précis:** We collected brain scans of 27 youth with classical CAH and 35 healthy controls. Portions of the prefrontal cortex, amygdala, and hippocampus were smaller in CAH youth compared to controls.

**Disclosure Summary:** MEG receives grant support from Novo Nordisk; consultant fees from Daiichi Sankyo, Ferring, Novo Nordisk, Nutrition & Growth Solutions, Pfizer, Sandoz, and Spruce Biosciences; serves on data safety monitoring boards for Ascendis, Millendo, and Tolmar; and receives royalties from McGraw-Hill and UpToDate.

## INTRODUCTION

Classical Congenital Adrenal Hyperplasia (CAH) is most commonly due to a mutation in the *CYP21A2* gene, affecting 1 in 15,000 live births, and is the most common cause of primary adrenal insufficiency in children (1). Inadequate production of cortisol and aldosterone, and overproduction of adrenal androgens, starting in the first trimester *in utero* can manifest in female newborns with the severe, classical forms of CAH as masculinized external genitalia (2,3). Given widespread expression of androgen and glucocorticoid receptors throughout the brain (4,5), there has been interest in understanding how hormonal imbalances related to CAH may impact distinct subregions of the developing brain (6). Several studies have begun to examine brain and behavioral alterations associated with CAH, with reported differences in emotional and memory processes in CAH patients. These include moderate-to-large reductions in short-term and working memory, which involve the hippocampus (7,8). Utilizing neuroimaging, abnormal amygdala volumes have been reported in children with CAH (9), with affected females exhibiting greater amygdala activity during functional magnetic resonance imaging (fMRI) emotional tasks (10) compared to control females (11). Adolescents with CAH also differ in their aversive ratings of fearful faces (11), exhibit poorer memory for negatively valenced stimuli on fMRI as compared to controls (10,12), and have impaired motivational inhibition on a reward-based anti-saccade task (13). Higher rates of anxiety disorders are also reported in youth with CAH compared to nationwide rates in healthy or other chronically ill pediatric populations (14). More recently, women (18-50 years) affected by CAH due to 21-hydroxylase deficiency were also found to have reduced hippocampal volumes and impaired cognitive performance on working memory and processing speed tests (15).

Both the amygdala and hippocampus are comprised of heterogenous cell types that can be categorized into subregions with distinct cytoarchitecture, with previous studies showing functional differences between the basal and lateral nuclei (which process high-level sensory input and emotional regulation) (16,17) and the central and basolateral nuclei (which are involved in reward learning and food intake) (18). Similarly, both structural and functional MRI studies have reported that hippocampal subfields, including the CA1, CA3, and dentate gyrus (DG), may have different involvement in learning and memory processes (19,20). The amygdala and hippocampus have both efferent and afferent connectivity with the prefrontal cortex (PFC), ultimately allowing for successful emotional regulation and learning and memory behaviors (21,22). However, it has yet to be studied how the distinct subregions of the amygdala and hippocampus are affected in patients with CAH.

The goal of the current study was to utilize state-of-the-art, high-resolution neuroimaging techniques to more fully characterize gray matter morphometry of the PFC, hippocampus, and amygdala in youth with classical CAH, compared to age- and sex-matched controls. CAH provides the opportunity to further understand how alterations in prenatal androgen and cortisol deficiency, as well as postnatal androgen and glucocorticoid exposure, may impact the developing brain. By combining 3D high-resolution T1- and T2-weighted structural sequences, there is substantial improvement of structural MRI to capture tissue contrast, which subsequently allows for enhanced segmentation of the amygdala into eight distinct subnuclei [lateral nucleus, basal nucleus, accessory basal nucleus, anterior amygdaloid area, medial nucleus, cortical nucleus, cortico-amygdaloid transition, and paralaminar nucleus] (23) and the hippocampus into six subfields [parasubiculum, presubiculum, subiculum, CA1, CA3, and DG] (24).

## PARTICIPANTS AND METHODS

### Study Participants

The study was cross-sectional and approved by the Institutional Review Board of the University of Southern California (USC) and Children’s Hospital Los Angeles (CHLA). Written consent was obtained from all parents or legal guardians, and/or participants, and all minors up to 14 years of age gave assent, in accordance with The Code of Ethics of the World Medical Association. Participants were recruited via flyers posted at CHLA and Keck School of Medicine of USC, with CAH participants recruited from the CHLA CAH Comprehensive Care Center. Health-related exclusionary criteria for all participants included prenatal drug or alcohol exposure, premature birth, serious medical illness (other than CAH), eating disorders, or psychotropic medication. Participants were screened for any significant neurological conditions (*e.g.*, epilepsy and traumatic head injury) and psychiatric/developmental disorders (*e.g.*, autism, attention deficit hyperactive disorder, schizophrenia, and self-harm tendencies) which, if present, barred participation. Participants were also screened for any factors that would prevent proper and safe usage of MRI, such as irremovable ferrous materials (*e.g.*, braces), uncorrectable vision impairments (*e.g.*, blind spots and colorblindness), need for hearing aids, or claustrophobia.

We studied a total of 62 children and adolescents between the ages of 8 and 18 years old at the time of their visit (Table 1), including 27 participants with classical CAH and 35 age- and sex-matched healthy controls with no significant medical conditions. Youth with CAH had either the salt-wasting (n=25) or simple-virilizing form (n=2), as diagnosed by positive newborn screen (n=12), or biochemically and/or by genotype (n=15; age of testing 11.2 ± 27.4 months). At the time of the study visit, patients with CAH were on daily glucocorticoid dosing (16.5 ± 4.7 mg/m^2^/day) with glucocorticoid dose equivalencies calculated based on growth-suppressing effects of longer-acting glucocorticoids compared to hydrocortisone (prednisone dose was multiplied by 5 and dexamethasone dose was multiplied by 80) (25). Almost all patients (n=26) were also treated with fludrocortisone (0.11 ± 0.04 mg/day).

**Table 1.**
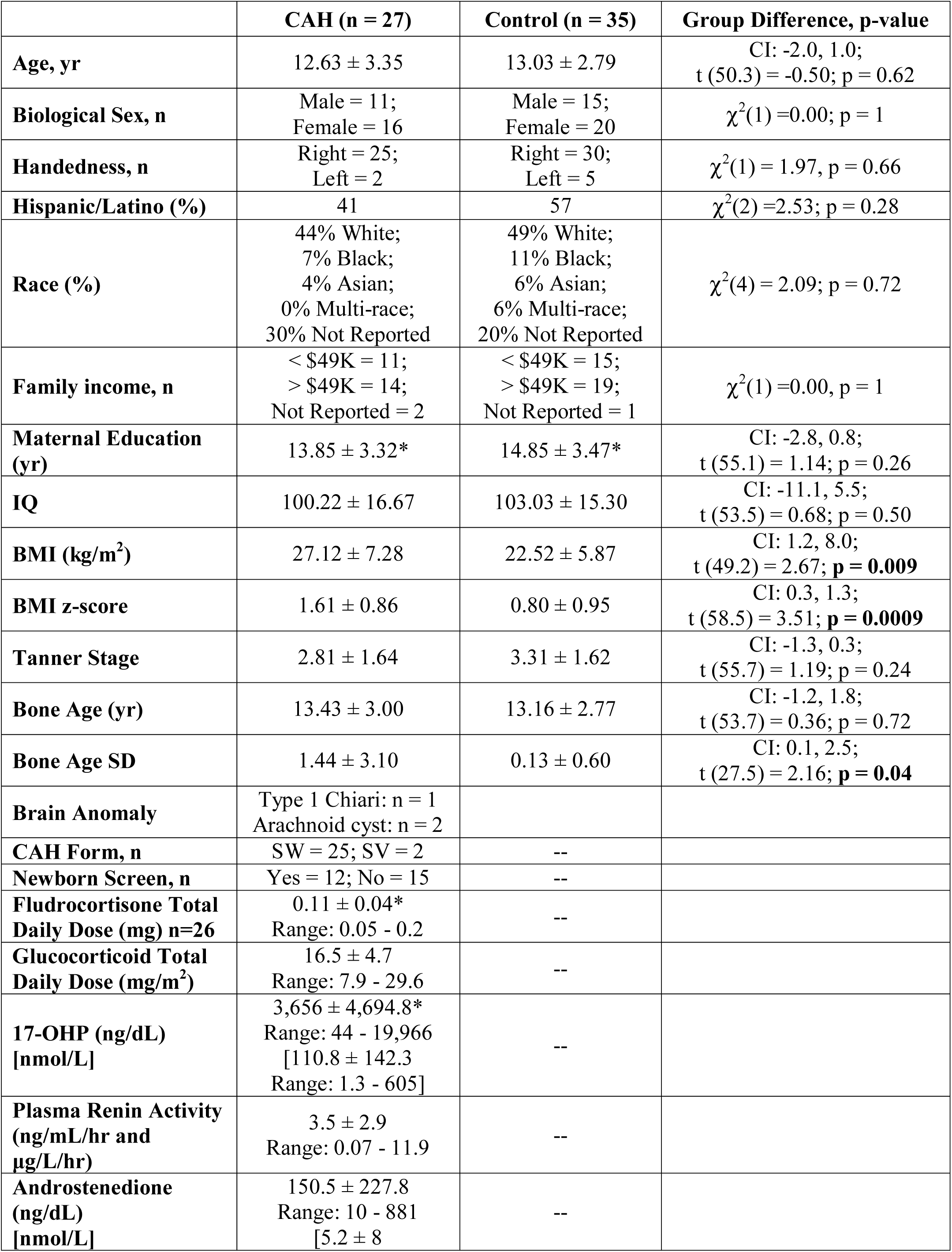
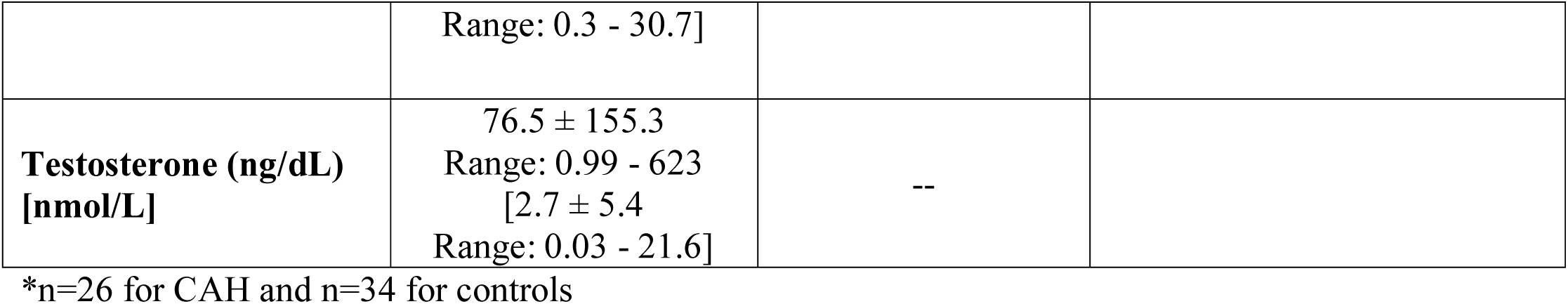
Study Participant Characteristics for CAH and Control Youth.

Anthropometric measures of height (cm) and weight (kg) were obtained in all participants. Pubertal Tanner staging was assessed by a pediatric endocrinologist. Body mass index (BMI) and BMI z-score were calculated by SAS based on 2000 Center for Disease Control Growth Chart data (26,27). A radiograph of the left hand was used to determine bone age (BA) using the Greulich-Pyle method (28) and read in a blinded fashion by a single pediatric endocrinologist (M.S.K.). BA SD was determined utilizing digital software (29). BA was obtained at the time of the study visit, or within several months of the visit if taken for clinical purposes. The individuals who had completed growth at the time of the study visit had their BA x-rays reviewed for earlier full maturity as adolescents.

There were no significant differences between the groups in terms of handedness, ethnicity/race composition, family income, maternal education, or IQ, as assessed by the Wechsler Adult Intelligence Scale (WASI) IV two-subtest test (30). Pubertal development was not significantly different between the two groups, although patients with CAH had higher BMI and BMI z-scores, as well as BA SD for their chronological age, as compared to control youth. After an overnight fast (12 h), and prior to routine morning medications in CAH youth, all participants had had their blood drawn at the CHLA Clinical Trials Unit for measurement of analytes including: 17-hydroxyprogesterone, androstenedione, plasma renin activity, and total testosterone by liquid chromatography tandem mass spectrometry (Quest Diagnostics Nichols Institute, San Juan Capistrano, CA).

### Magnetic Resonance Imaging (MRI) Acquisition

All images were collected on a Siemens Magnetom Prisma 3 Tesla MRI scanner using a 32-channel head coil at University of Southern California’s Center for Image Acquisition. T1-weighted structural imaging was acquired using a sagittal whole brain MPRAGE sequence (TR = 2400 ms, TE = 2.22 ms, flip angle = 8°, BW = 220 Hz/Px, FoV = 256 mm, 208 slices, and 0.8-mm isotropic voxels, with a GRAPPA phase-encoding acceleration factor of 2). T2-weighted variable flip angle turbo spin-echo sequence was also collected (TR = 3200 ms, TE = 563 ms, BW = 744 Hz/Px, FoV = 256 mm, 208 slices, 0.8-mm isotropic voxels, and 3.52-ms echospacing, with a GRAPPA phase-encoding acceleration factor of 2). Anterior-posterior and posterior-anterior spin echo field maps were also obtained (TR = 8000 ms, TE = 66.0 ms, flip angle = 90°, BW = 2290 Hz/Px, FoV = 208 mm, 72 slices, and 2.0-mm isotropic voxels, with a multi-band acceleration factor of 1). A radiologist reviewed all scans for incidental findings of gross abnormalities.

### MRI Analysis

#### Whole Brain Segmentation

Structural image processing, including whole brain segmentation with automated labeling of different neuroanatomical structures, was performed using FreeSurfer v6.0 (http://surfer.nmr.mgh.harvard.edu) (31,32). Standard quality control procedures were as follows: 1) all raw images were visually inspected for motion prior to processing and 2) post-processed images were visually inspected by a trained operator for accuracy of segmentation for each scan per participant (33). No manual intervention (*i.e.*, subcortical editing) was performed. In addition to total gray matter, cerebrospinal fluid (CSF) and intracranial (ICV) volume were extracted as well as five *a priori* prefrontal regions of interest (ROI) using the Desikan-Killiany Atlas, including the superior, rostral middle, caudal middle, lateral orbitofrontal, and medial orbitofrontal regions for both the right and left hemisphere.

#### Amygdala and Hippocampal Segmentation

Details of the *in vivo* amygdala probabilistic atlas construction, validation, estimates of individual differences, and comparison with previous atlases have been previously published (23). Each participant’s image was registered to the CIT168 atlas using a B-spline bivariate symmetric normalization diffeomorphic registration algorithm from ANTs (34). Implementation of the inverse diffeomorphism resulted in a probabilistic segmentation of each participant’s left and right total amygdala estimates, as well as the following eight distinct bilateral regions of interest (ROI): lateral nucleus (lral nucleus (LA)), dorsal and intermediate divisions of the basolateral nucleus (BLDI), ventral division of the basolateral nucleus and paralaminar nucleus (BLVPL), basomedial nucleus (BM), central nucleus (CEN), cortical and medial nuclei (CMN), amygdala transition areas (ATA), amygdalostriatal transition area (ASTA), and anterior amygdala area (AAA). In the creation and validation of the CIT168 atlas, Tyszka and Pauli (23) established that a contrast-to-noise ratio (CNR) > 1 provides a robust estimation of the ground truth volumes. Using their established formula, the CNR for our sample was 1.21 and 1.01 for the T1- and T2-weighted scans suggesting that we had sufficient CNR to establish amygdala boundaries using this method.

T1- and T2-weighted images for each participant were also utilized to quantify six hippocampal subfields, including the parasubiculum, presubiculum, subiculum, CA1, CA3, and dentate gyrus (DG) using the computational atlas (24) available in FreeSurfer v6.0 (31). This method provides hippocampal subfield volumetric measures that more closely align with histological measurements, compared to alternative automated segmentation algorithms and previous versions of the software (24).

In addition to absolute volumes of amygdala subnuclei and hippocampal subfields, we calculated a relative volume fraction (RVF) for each ROI by normalizing it to the respective total amygdala volume in each hemisphere (ROI probabilistic volume divided by total amygdala probabilistic volume). In addition to examining overall differences in volume, RVF allows for determining if the amygdala or hippocampal relative composition differed in youth affected with CAH as compared to controls (*e.g.*, testing if within the amygdala the proportions of each nuclei are relatively smaller or larger to other nuclei).

### Statistical Analyses

All data analyses were performed in Rstudio v1.2 (Boston, MA http://www.rstudio.com/) using linear multiple regression and linear-mixed models from package nlme v3.1 (https://CRAN.R-project.org/package=nlme). Given group differences in BMI z-scores, BA SD, and ICV (Tables 1 and 2), these variables were included as covariates in all subsequent analyses when applicable. Separate multiple regressions were first implemented to examine how group, sex, and their interaction (group*sex) predicted CSF and ICV, while controlling for BMI z-score and BA SD. In addition, similar models were performed for each *a priori* PFC and total amygdala and hippocampus volumes for the right and left hemispheres, while adjusting means for BMI z-score, BA SD, and ICV. To assess amygdala subnuclei and hippocampal subfields, separate multi-level models were then performed to investigate group (CAH vs control) and group-by-region effects for both the absolute volumes and RVF of the eight amygdala subnuclei and six hippocampal subfields across both hemispheres (right, left), with the within-subject variable, while also controlling for BMI z-score, BA SD, and ICV (for absolute volumes). Post-hoc tests were then performed to probe significant interactions (group-by-region), correcting for multiple comparisons using the Holm method. Lastly, for regions showing significant group differences in brain structure, a within-group assessment of relationships between brain structure and clinical features was performed in youth with CAH using multiple regression. These clinical features included: BA SD, testosterone and androstenedione levels, and glucocorticoid daily dose (mg/m^2^/day).

**Table 2.** Brain Volumes Mean and Standard Deviation for CAH and Control Youth.

|  | CAH |  | Controls |  | F-statistic, p-value |
| --- | --- | --- | --- | --- | --- |
|  | Males | Females | Males | Females |  |
| Total Brain Volumes |  |  |  |  |  |
| ICV | 1559479 ± 163595.50 | 1438723 ± 157480.10 | 1647836 ± 136404.50 | 1515500 ± 105509.60 | Group: F(1, 55) =6.34, **p = 0.01**, ηp² =0.10<br>Sex: F(1, 55) =10.7, **p = 0.002**, ηp² =0.17 |
| CSF | 1123.47 ± 445.98 | 1023.49 ± 187.06 | 911.70 ± 203.93 | 812.71 ± 145.31 | Group: F(1, 56) =6.46, **p = 0.01**, ηp² =0.10<br>Sex: F(1, 56) =2.1, p = 0.16, ηp² =0.04 |
| Total Gray Matter | 605884.30 ± 58869.03 | 541305.40 ± 55596.74 | 645383.50 ± 39360.51 | 584338.20 ± 40641.82 | Group: F(1, 55) =1.48, p = 0.23, ηp² =0.03<br>Sex: F(1, 55) =10.9, **p = 0.002**, ηp² =0.17 |
| Amygdala |  |  |  |  |  |
| Right | 1724.07 ± 172.76 | 1557.87 ± 212.42 | 1873.87 ± 191.82 | 1689.65 ± 215.5 | Group: F(1, 55) =1.10, p = 0.10, ηp² =0.05<br>Sex: F(1, 55) =1.47, p = 0.23, ηp² =0.03 |
| Left | 1516.58 ± 166.03 | 1437.96 ± 216.11 | 1658.45 ± 176.89 | 1551.60 ± 176.05 | Group: F(1, 55) =1.10, p = 0.30, ηp² =0.02<br>Sex: F(1, 55) =0.0004, p = 0.98, ηp² <0.0001 |
| Hippocampus |  |  |  |  |  |
| Right | 4069.85 ± 817.28 | 3843.82 ± 390.82 | 4288.53 ± 337.15 | 4136.89 ± 407.37 | Group: F(1, 55) =2.00, p = 0.16, ηp² =0.04<br>Sex: F(1, 55) =1.72, p = 0.20, ηp² =0.03 |
| Left | 3750.32 ± 323.39 | 3662.90 ± 389.04 | 4149.08 ± 378.19 | 3921.53 ± 362.23 | Group: F(1, 55) =6.97, **p = 0.01**, ηp² =0.11<br>Sex: F(1, 55) =0.94, p = 0.34, ηp² =0.02 |
| Superior Frontal |  |  |  |  |  |
| Right | 35709.82 ± 4249.97 | 31901.44 ± 4740.10 | 39502.47 ± 3196.66 | 35363.80 ± 3014.59 | Group: F(1, 55) =5.66, **p = 0.02**, ηp² = 0.09<br>Sex: F(1, 55) =4.01, **p = 0.05**, ηp² = 0.07 |
| Left | 32669.64 ± 2806.22 | 29490.69 ± 4125.43 | 35863.13 ± 4071.86 | 32581.2 ± 2958.53 | Group: F(1, 55) =5.66, **p = 0.02**, ηp² = 0.09<br>Sex: F(1, 55) =4.01, **p = 0.05**, ηp² = 0.07 |
| Caudal Middle Frontal |  |  |  |  |  |
| Right | 7700.45 ± 1681.72 | 7104.75 ± 1613 | 8634.20 ± 1796.09 | 8832.15 ± 1426.20 | Group: F(1, 55) =8.46, **p = 0.005**, ηp² =0.13<br>Sex: F(1, 55) =2.92, p = 0.09, ηp² = 0.05 |
| Left | 8221.09 ± 1427.20 | 7382.88 ± 1787.81 | 9615.33 ± 1707.55 | 8505.9 ± 1402.16 | Group: F(1, 55) =5.31, **p = 0.02**, ηp² =0.09<br>Sex: F(1, 55) =1.44, p = 0.24, ηp² = 0.03 |
| Rostral Middle Frontal |  |  |  |  |  |
| Right | 15967.36 ± 2311.01 | 13749.25 ± 1837.47 | 16691.80 ± 1832.05 | 15089.20 ± 2119.11 | Group: F(1, 55) =0.34, p = 0.56, ηp² =0.006<br>Sex: F(1, 55) =2.81, p = 0.09, ηp² = 0.05 |
| Left | 15490.27 ± 1944.1 | 13004.19 ± 2068.18 | 16508 ± 2409.9 | 15561.95 ± 2010.04 | Group: F(1, 55) =7.18, **p = 0.009**, ηp² =0.12<br>Sex: F(1, 55) =0.32, p = 0.58, ηp² = 0.006 |
| Medial Orbitofrontal |  |  |  |  |  |
| Right | 5207.55 ± 694.62 | 5035.12 ± 794.31 | 5218.93 ± 920.63 | 5013 ± 800.24 | Group: F(1, 55) =2.37, p = 0.13, ηp² = 0.04<br>Sex: F(1, 55) =0.91, p = 0.34, ηp² = 0.02 |
| Left | 5194.27 ± 1266.51 | 5030.06 ± 563.45 | 5319.87 ± 661.67 | 4888.75 ± 641.86 | Group: F(1, 55) =1.39, p = 0.24, ηp² = 0.02<br>Sex: F(1, 55) =0.04, p = 0.84, ηp² = 0.0007 |
| Lateral Orbitofrontal |  |  |  |  |  |
| Right | 11313.73 ± 1017.28 | 10217.62 ± 1261.01 | 11351 ± 1243.4 | 10405.1 ± 868.41 | Group: F(1, 55) =3.28, p = 0.07, ηp² = 0.06<br>Sex: F(1, 55) =2.43, p = 0.12, ηp² = 0.04 |
| Left | 11154.36 ± 1196.43 | 10356.44 ± 1168.65 | 11593 ± 1159.35 | 10690.65 ± 969.28 | Group: F(1, 55) =0.07, p = 0.79, ηp² =0.001<br>Sex: F(1, 55) =0.61, p = 0.43, ηp² = 0.01 |

ICV (intracranial volume), CSF (cerebrospinal fluid)

## RESULTS

### Central Nervous System Abnormalities

During radiology review, one CAH patient was found to have a Type 1 Chiari malformation, and two CAH patients were found to have arachnoid cysts, with one cyst located near the temporal lobe and the other near the cerebellum. Because central nervous system abnormalities have been previously reported in patients with CAH (15), analyses were performed including these participants; however, follow-up analyses were also examined excluding the data from these three patients to ensure they were not driving group effects. Sample characteristics were similar between the groups when excluding the three CAH patients with brain anomalies.

### Total Brain Volumes

A significant main effect was seen for both ICV and CSF, with youth with CAH having smaller ICV, but larger CSF volumes, compared to controls (Table 2, Figure 1A and 1B). No significant differences were seen in overall cortical gray matter (Figure 1C). For the medial temporal lobe volumes, total amygdala and hippocampus volumes were smaller on average in CAH youth compared to controls, although in statistical testing only the total left hippocampus volume was significantly smaller in CAH (p = 0.01). (Table 2, Figure 2A and B). For the PFC region, the bilateral superior frontal, bilateral caudal middle frontal, and left rostral middle frontal regions were significantly smaller in CAH youth compared to controls (Table 2, Figure 2C and D). A trend was also seen in smaller right lateral orbitofrontal volumes in CAH youth compared to controls, which did not reach statistical significance (p = 0.07). Group-by-sex effects were not significant in any of the models (data not shown).

**Figure 1.**
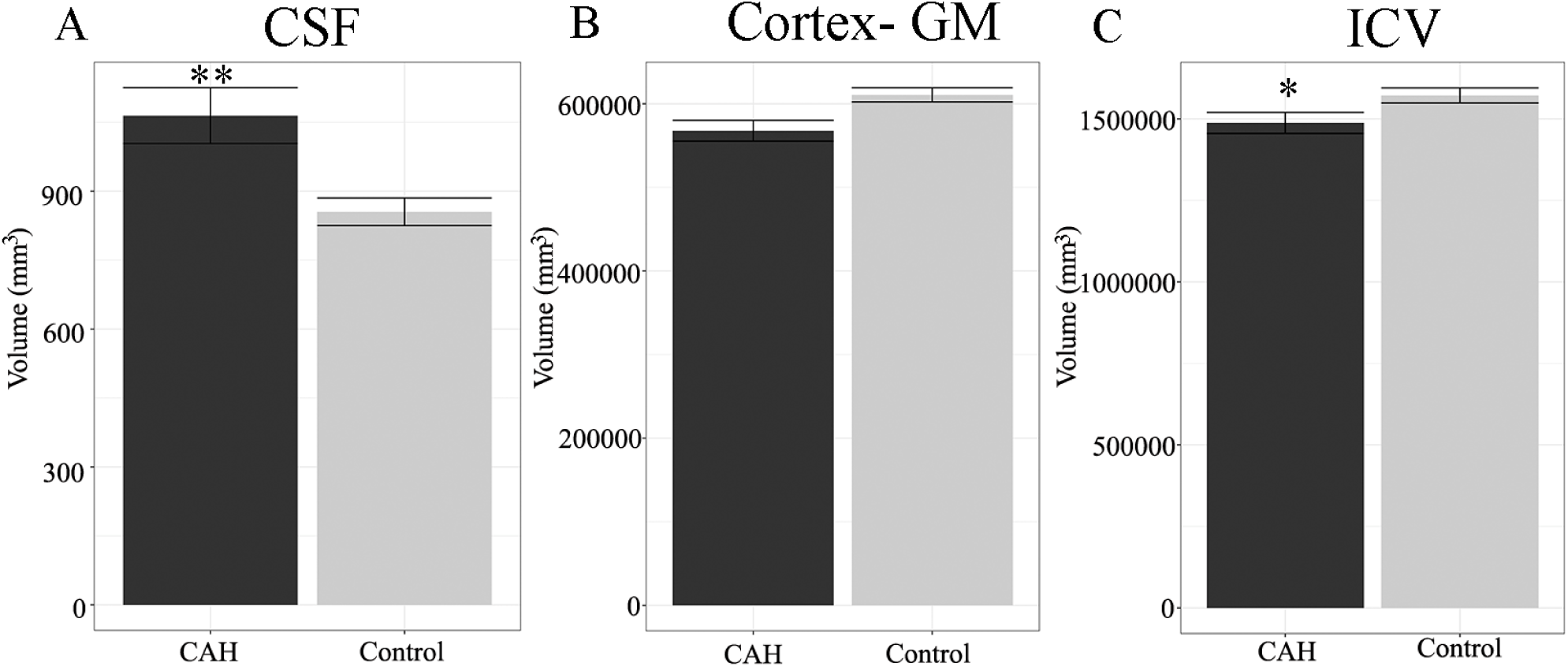
Global brain volumes in youth with classical CAH and age- and sex-matched healthy controls. Plots reflect means and standard error of volumes for: A) Cerebrospinal fluid (CSF), B) Cortex, gray matter (GM), and C) Intracranial volume (ICV). A) CSF is significantly larger in CAH compared to control youth (** p ≤ 0.01). C) ICV is significantly smaller in CAH compared to control youth (** p ≤ 0.01).

**Figure 2.**
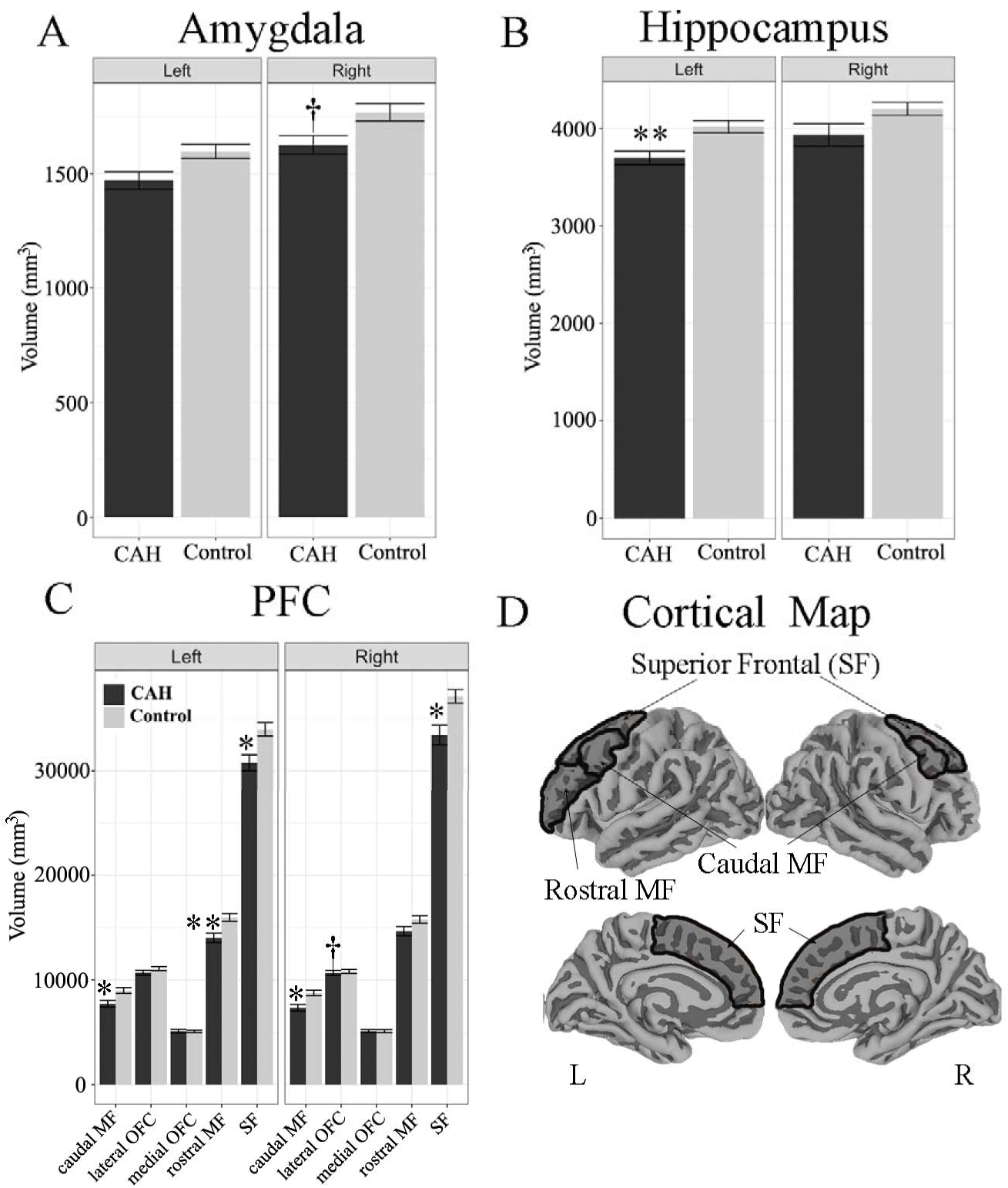
Total amygdala, hippocampus, and prefrontal cortex (PFC) volumes in youth with classical CAH and controls by hemisphere. Plots reflect means and standard error of volumes for: A) Amygdala, B) Hippocampus, and C) Prefontal cortex (PFC) regions of interest. A) A trend level difference was seen for smaller total right amygdala volumes in CAH compared to control youth (^†^p = 0.10). B) Left hippocampus is significantly smaller in CAH compared to control youth (** p ≤ 0.01). C) Bilateral superior frontal (SF) cortex volumes (* p ≤ 0.05) and caudal middle frontal (MF) cortex volumes (* p ≤ 0.05) were significantly smaller in CAH compared to control youth. Left rostral middle frontal volumes were significantly smaller in CAH compared to control youth (** p ≤ 0.01). A trend level difference was seen for smaller right lateral orbitofrontal cortex (OFC) volumes in CAH compared to control youth (^†^p = 0.07). D) Significant PFC regions of interest mapped to the cortex for visualization (dark gray).

### Amygdala Subnuclei and Hippocampal Subfields

Significant group [F(1,55) = 11.70, p = 0.001, ηp^2^ = 0.006] and group-by-region interactions [F(8,1020) = 5.64, p < 0.0001, ηp^2^ = 0.03] were seen in the absolute volumes for amygdala subnuclei (Figure 3A). Follow-up post-hoc analyses revealed that the largest amygdala subnucleus, the LA, was significantly smaller in CAH patients as compared to controls (Table 3, Figure 3A). No main effect of group [F(1,55) = 0.08, p = 0.77, ηp^2^ < 0.001] or group-by-region interaction [F(8,1020) = 1.16, p = 0.32, ηp^2^ = 0.008] was seen for amygdala RVF (Table 3, Figure 3A).

**Table 3.** Amygdala Nuclei Mean and Standard Deviation for CAH and Control Youth

|  | CAH (n=26) |  | Controls (n=35) |  | Post-hoc Test of Group Difference |
| --- | --- | --- | --- | --- | --- |
|  | Total | RVF | Total | RVF | Total |
| Lateral Nucleus (LA) |  |  |  |  |  |
| Right | 307.22 ± 40.84 | 0.157 ± 0.013 | 331.86 ± 46.67 | 0.159 ± 0.012 | Means<sub>adj</sub> = -18.86,<br>χ<sup>2</sup>(1) = 24.88;<br>**p < 0.0001** |
| Left | 297.32 ± 42.92 | 0.157 ± 0.012 | 318.99 ± 46.3 | 0.158 ± 0.013 |  |
| Basolateral Nucleus (BLDI) |  |  |  |  |  |
| Right | 192.26 ± 22.75 | 0.098 ± 0.004 | 202.12 ± 24.55 | 0.097 ± 0.004 | Means<sub>adj</sub> = -6.46,<br>χ<sup>2</sup>(1) = 2.92; p = 0.08 |
| Left | 187.24 ± 22.87 | 0.099 ± 0.005 | 198.89 ± 23.25 | 0.099 ± 0.004 |  |
| Basomedial Nucleus (BM) |  |  |  |  |  |
| Right | 109.04 ± 17.47 | 0.056 ± 0.005 | 114.37 ± 15.76 | 0.055 ± 0.004 | Means<sub>adj</sub> = -.79,<br>χ<sup>2</sup>(1) = 0.44; p = 0.83 |
| Left | 106.64 ± 14.14 | 0.057 ± 0.004 | 111.48 ± 13.88 | 0.056 ± 0.006 |  |
| Central Nucleus (CEN) |  |  |  |  |  |
| Right | 46.95 ± 8.06 | 0.024 ± 0.003 | 49.04 ± 6.95 | 0.024 ± 0.002 | Means<sub>adj</sub> = 2.16,<br>χ<sup>2</sup>(1) = 0.33; p = 0.57 |
| Left | 46.28 ± 7.53 | 0.025 ± 0.003 | 48.46 ± 6.79 | 0.025 ± 0.003 |  |
| Cortical and Medial Nuclei (CMN) |  |  |  |  |  |
| Right | 151.04 ± 22.24 | 0.077 ± 0.006 | 156.99 ± 19.61 | 0.078 ± 0.007 | Means<sub>adj</sub> = -3.11,<br>χ<sup>2</sup>(1) = 0.67; p = 0.41 |
| Left | 147.21 ± 19.26 | 0.078 ± 0.006 | 156.07 ± 19.44 | 0.076 ± 0.007 |  |
| Ventral Division of BLDI and Paralaminar Nucleus (BLVPL) |  |  |  |  |  |
| Right | 112.21 ± 13.2 | 0.058 ± 0.005 | 120.63 ± 16.43 | 0.058 ± 0.004 | Means<sub>adj</sub> = -4.39,<br>χ<sup>2</sup>(1) = 1.35; p = 0.25 |
| Left | 111.06 ± 14.55 | 0.059 ± 0.004 | 120 ± 15.87 | 0.060 ± 0.004 |  |
| Amygdala Transition Areas (ATA) |  |  |  |  |  |
| Right | 69.98 ± 10.49 | 0.036 ± 0.004 | 75.44 ± 12.05 | 0.037 ± 0.004 | Means<sub>adj</sub> = -0.60,<br>χ<sup>2</sup>(1) = 0.25; p = 0.87 |
| Left | 67.7 ± 10.61 | 0.036 ± 0.003 | 69.63 ± 9.21 | 0.035 ± 0.003 |  |
| Amygdalostratial Transition Area (ASTA) |  |  |  |  |  |
| Right | 63.48 ± 8.1 | 0.033 ± 0.004 | 69.99 ± 9.4 | 0.034 ± 0.004 | Means<sub>adj</sub> = -2.18,<br>χ<sup>2</sup>(1) = 0.33; p = 0.56 |
| Left | 63.12 ± 8.6 | 0.034 ± 0.004 | 69.56 ± 10.13 | 0.035 ± 0.004 |  |
| Anterior Amygdala Area (AAA) |  |  |  |  |  |
| Right | 61.74 ± 7.92 | 0.032 ± 0.003 | 61.44 ± 7.49 | 0.030 ± 0.003 | Means<sub>adj</sub> = 3.64,<br>χ<sup>2</sup>(1) = 0.92; p = 0.34 |
| Left | 59.14 ± 6.84 | 0.032 ± 0.003 | 60.76 ± 7.66 | 0.031 ± 0.004 |  |

**Figure 3.**
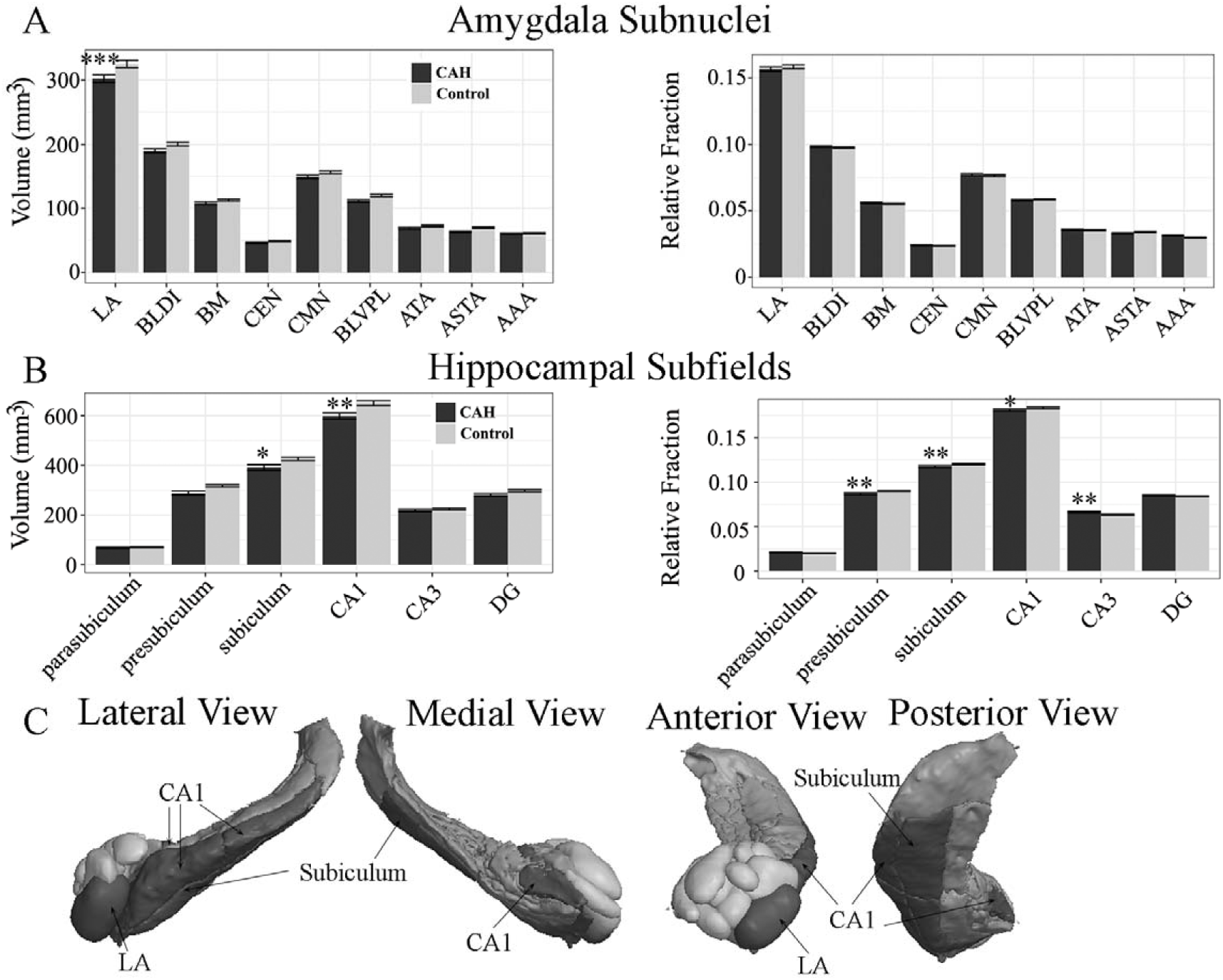
Amygdala subnuclei and hippocampal subfield differences in youth with classical CAH and controls. Means and standard errors plotted for absolute (left) and relative volume fraction (right) across: A) Eight amygdala subnuclei, and B) Six hippocampal subfields. A) Absolute lateral nucleus of the amygdala (LA) volumes were significantly smaller in CAH as compared to control youth (*** p ≤ 0.001). B) Absolute subiculum and CA1 volumes were significantly smaller in CAH compared to control youth (*p ≤ 0.05 and ** p ≤ 0.01, respectively). Relative volume fractions were also significantly smaller for the presubiculum (** p ≤ 0.01), subiculum (** p ≤ 0.01), and CA1 (* p ≤ 0.05) in CAH compared to control youth. Relative volume fractions were also significantly larger for the CA3 subfield (** p ≤ 0.01). C) 3D rendering of amygdala (light gray) and hippocampal regions (dark gray) on a representative CAH participant with significantly smaller absolute volumes per region in CAH compared to control youth (black).

Significant group [F(1,56) = 8.99, p = 0.004, ηp^2^ = 0.02] and group-by-region interactions [F(5,671) = 7.80, p < 0.0001, ηp^2^ = 0.03] were also seen in the absolute volumes for the hippocampal subfields (Figure 3B). Follow-up post-hoc analyses revealed that the subiculum and CA1 subfields were significantly smaller in CAH youth compared to controls (Figure 3B). For hippocampal RVF, the overall main effect of group was not significant [F(1,57) = 1.82, p = 0.18, ηp^2^ = 0.002], but the group-by-region interaction was significant [F(6,671) = 5.07, p < 0.0001, ηp^2^ = 0.03] (Figure 3B). Even after accounting for smaller hippocampal volumes, the presubiculum, subiculum, and CA1 were proportionately smaller, whereas CA3 was proportionately larger, relative to overall hippocampal volumes in CAH patients as compared to controls.

### Brain Volumes and Clinical Features in CAH Youth

No associations were seen between brain volumes and CAH clinical features (data not shown), including markers of androgen excess (*e.g.*, BA SD), total testosterone and androstenedione levels, 17-hydroxyprogesterone, or glucocorticoid daily dose (Table 1).

Finally, sensitivity analyses were performed for significant findings excluding the three CAH patients with brain anomalies. Results were nearly identical, although some regions, including the total left hippocampus, bilateral superior frontal cortex, and the relative volume fraction of CA1, became trend-level, and the left caudal middle frontal cortex was no longer significant (Table 5).

**Table 4.** Hippocampal Subfields Mean and Standard Deviation for CAH and Control Youth

|  | CAH<br>(n = 27) |  | Controls<br>(n = 35) |  | Post-hoc Test of Group by Region |  |
| --- | --- | --- | --- | --- | --- | --- |
|  | Total | RVF | Total | RVF | Total | RVF |
| Parasubiculum |  |  |  |  |  |  |
| Right | 69.15 ± 16.08 | 0.021 ± 0.004 | 70.27 ± 14.83 | 0.020 ± 0.004 | Means<sub>adj</sub> = 7.76,<br>χ<sup>2</sup>(1) = 0.47;<br>p = 0.49 | Means<sub>adj</sub> = -0.0008,<br>χ<sup>2</sup>(1)=0.49;<br>p = 0.48 |
| Left | 68.49 ± 12.81 | 0.022 ± 0.004 | 70.23 ± 12.14 | 0.021 ± 0.004 |  |  |
| Presubiculum |  |  |  |  |  |  |
| Right | 291.48 ± 67.89 | 0.085 ± 0.007 | 319.14 ± 36.62 | 0.089 ± 0.006 | Means<sub>adj</sub> = -20.67,<br>χ<sup>2</sup>(1) = 3.36;<br>p = 0.07 | Means<sub>adj</sub> = -0.003,<br>χ<sup>2</sup>(1) = 7.99;<br>**p = 0.005** |
| Left | 285.38 ± 47.4 | 0.089 ± 0.009 | 317.45 ± 36.61 | 0.092 ± 0.005 |  |  |
| Subiculum |  |  |  |  |  |  |
| Right | 401.79 ± 111.56 | 0.117 ± 0.01 | 429.61 ± 55.35 | 0.120 ± 0.007 | Means<sub>adj</sub> = -25.76,<br>χ<sup>2</sup>(1) = 5.22;<br>**p = 0.02** | Means<sub>adj</sub> = -0.003,<br>χ<sup>2</sup>(1) = 6.29;<br>**p = 0.01** |
| Left | 381.15 ± 51.69 | 0.119 ± 0.007 | 423.23 ± 54.67 | 0.122 ± 0.007 |  |  |
| CA1 |  |  |  |  |  |  |
| Right | 626.94 ± 107.6 | 0.184 ± 0.009 | 666.49 ± 87.09 | 0.185 ± 0.011 | Means<sub>adj</sub> = -42.40,<br>χ<sup>2</sup>(1) = 14.13;<br>**p < 0.0002** | Means<sub>adj</sub> = -0.0025,<br>χ<sup>2</sup>(1)=4.89;<br>**p = 0.03** |
| Left | 571.1 ± 61.85 | 0.179 ± 0.008 | 634.74 ± 82.84 | 0.183 ± 0.009 |  |  |
| CA3 |  |  |  |  |  |  |
| Right | 227.15 ± 39.9 | 0.067 ± 0.007 | 229.55 ± 32.18 | 0.064 ± 0.006 | Means<sub>adj</sub> = 4.05,<br>χ<sup>2</sup>(1) = 0.13;<br>p = 0.71 | Means<sub>adj</sub> = 0.003,<br>χ<sup>2</sup>(1) = 6.25;<br>**p = 0.01** |
| Left | 209.96 ± 24.22 | 0.066 ± 0.006 | 217.84 ± 29.65 | 0.063 ± 0.006 |  |  |
| Dentate Gyrus (DG) |  |  |  |  |  |  |
| Right | 289.33 ± 48.44 | 0.085 ± 0.005 | 304.37 ± 38.47 | 0.085 ± 0.005 | Means<sub>adj</sub> = -8.09,<br>χ<sup>2</sup>(1) = 0.51;<br>p = 0.47 | Means<sub>adj</sub> = 0.0008,<br>χ<sup>2</sup>(1) = 0.46;<br>p = 0.50 |
| Left | 272.55 ± 27.73 | 0.086 ± 0.004 | 292.08 ± 35.31 | 0.084 ± 0.004 |  |  |

**Table 5.** Sensitivity Analyses Excluding Three CAH Patients with Brain Anomalies

| **Brain Region** | **Group**<br>**Difference**<br>**p-value** |
| --- | --- |
| Intracranial Volume (ICV) | **0.008** |
| Cerebrospinal Fluid (CSF) | **0.007** |
| Left Hippocampus | 0.07 |
| Right Superior Frontal | 0.08 |
| Left Superior Frontal | 0.09 |
| Right Caudal Middle Frontal | **0.01** |
| Left Caudal Middle Frontal | 0.11 |
| Left Rostral Middle Frontal | **0.02** |
| Lateral Nucleus (LA) | **< 0.001** |
| Subiculum | **0.001** |
| CA1 | **< 0.001** |
| Presubiculum Relative Volume Fraction (RVF) | **0.001** |
| Subiculum Relative Volume Fraction (RVF) | **0.003** |
| CA1 Relative Volume Fraction (RVF) | 0.07 |

## DISCUSSION

Our study quantified regional differences in gray matter morphometry of the prefrontal cortex, hippocampus, and amygdala in youth affected with classical CAH and age- and sex-matched controls. We found smaller total volumes of the amygdala and hippocampus in CAH youth, similar to prior findings (15,35), and expanded upon these findings to show regional volume differences in the prefrontal cortex (smaller volumes of the superior and caudal middle frontal cortex), lateral nucleus of the amygdala, and subiculum and CA1 subregions of the hippocampus in CAH youth compared to controls. We also found CAH youth to have smaller intracranial volumes and increased cerebral spinal fluid, signifying whole brain atrophy as well as regional specific alterations in the prefrontal and medial temporal lobe regions during childhood and adolescent development.

Over the last two decades, excluding various case studies (36-38), only a small number of experimental MRI studies have examined gray matter volumes in children and adults affected with CAH. The first study included 39 patients (3 months to 26 years old), 11 of whom displayed temporal lobe atrophy on T1- and T2-weighted MRI; albeit the study design did not include a true control group (39). Another study quantified total amygdala and hippocampal volumes in children with CAH as compared to controls, showing smaller amygdala volumes but no difference in hippocampal volumes, in both males and females with CAH (35). These findings are in contrast, however, to a more recent study that found only a trend-level difference in the left amygdala and a significantly smaller right hippocampus in 19 adult women with CAH compared to unaffected women (15). Of note, the latter two studies also reported global level differences in brain structure, including trend-level differences in total cerebral volume in female youth with CAH (35) and increased CSF in adult women with CAH (15). Our findings suggest both smaller intracranial volumes, as well as increased CSF, in both males and females with CAH using a higher-resolution MRI scan and both T1- and T2-weighted images to improve accuracy in quantifying gray and white matter boundaries. Although we found that, in general, total bilateral amygdala and hippocampal volumes were smaller in CAH compared to matched controls, the differences in absolute amygdala and hippocampal volumes between CAH and control youth may be a consequence of overall smaller brain volumes given that group differences exist in ICV. After accounting for smaller intracranial volumes in our final models, only the left hippocampus was significantly smaller in both males and females with CAH compared to control youth, suggesting that differences in the amygdala may be a function of global differences in brain volumes, whereas the smaller left hippocampal volume is not a function of the overall difference in brain size.

Our interest in examining the PFC, amygdala, and hippocampus in CAH stems from the high concentrations of androgens, as well as mineralocorticoid and glucocorticoid receptors, in these neural regions (4,40-42). However, both the amygdala and the hippocampal formation have heterogeneous subregions in terms of their cytoarchitecture and projections (43,44). A major signaling pathway of the hippocampus, known as the perforant path, includes projections from the entorhinal cortex to both the DG and CA3, and then onto the CA1, hippocampal subfields. The DG is comprised of dense granule cells that send their projections, known as mossy fibers, to CA3, whereas the pyramidal cells in CA3 send their axons, known as the Schaffer collaterals, to CA1. The CA1 projects to the subiculum and entorhinal cortex, and is the main output of the hippocampus (44). Amygdala nuclei also have distinct connectivity patterns. The basolateral portion of the amygdala, including the lateral (LA region) and basal (BLDI, BM, and BLVPL regions) nuclei, receives sensory and regulatory information from the thalamus and PFC respectively. Within the amygdala, the basolateral nuclei send projections to the central and medial nuclei, which project to the hypothalamus and the bed nucleus of the stria terminalis (BNST) (43). Moreover, reciprocal connections exist between the hippocampus and amygdala, and the PFC and amygdala, with basal nuclei of the amygdala projecting to the entorhinal cortex and PFC, and subiculum and CA1 subfields of the hippocampus projecting back to the basolateral amygdala (45). Our findings suggest that within these heterogeneous medial temporal lobe structures, the lateral nucleus of the amygdala, and the subiculum and CA1 hippocampal subregions are the most affected in youth with CAH, which could have clinical consequences for various emotional and reward related behaviors.

Mechanistically, it remains unclear as to what feature(s) inherent to CAH are responsible for decreases in cortical volumes in the prefrontal and medial temporal lobes. CAH includes prenatal glucocorticoid deficiency, excess prenatal and postnatal androgens, and postnatal glucocorticoid treatment. Although clinical features have been found to relate to some cognitive tests or other brain biomarkers (15), we and others have not yet found markers of glucocorticoid or androgen exposure to predict amygdala or hippocampal volumes (15,35). Moreover, group differences did not significantly differ between the sexes (*i.e.*, group*sex interaction terms were not significant in any of our models), suggesting that both males and females with CAH are equally affected. In addition, well-known sex differences, in regard to females having smaller gray matter volumes compared to males, were preserved in youth with CAH. It is feasible that amygdala, hippocampus, and PFC volumes may be smaller as a result of their potentially diminished role in neural regulation of the hypothalamic-pituitary-adrenal (HPA) axis, as a result of primary adrenal insufficiency in CAH (46). Those with other pediatric clinical disorders associated with HPA dysfunction, such as hypercortisolism seen in Cushing syndrome, also have been shown to have smaller total brain volumes, including the amygdala and hippocampus. In addition, suppressed cortisol awakening responses have also been associated with smaller PFC and hippocampal volumes in individuals at high risk for psychosis based on desensitized HPA responsivity (47). Further support for this idea may stem from both absolute and proportionately smaller volumes localized to the subiculum and CA1 hippocampal subfields in youth with CAH, as the CA1 responds to glucocorticoids in a dose-response fashion and the subiculum plays a key role in inhibiting the HPA axis (48,49). Taken together, these data suggest that disruptions of HPA homeostasis may result in altered amygdala, hippocampus, and PFC volumes (50). Alternatively, a lack of association between brain volumes and exposures due to treatment or hormone concentrations may be muddled by individual differences in glucocorticoid-related genes, which may otherwise complicate hormone concentrations and clearance, as well as physiological neural responses (51). In addition, neurologic impairments, including possible brain injury, may result from adrenal crises associated with CAH which merits further study (7). More research is needed to better understand the biological mechanisms that may contribute to regional atrophy of the amygdala, hippocampus, and PFC in youth with CAH.

Strengths of the current study include the high-resolution T1- and T2-weighted MRI approach allowing for more accurate quantification of tissue as well as segmentation of the amygdala and hippocampus into distinct subregions. Although equal or larger than previous studies (15,35), our CAH sample size is still relatively small, which may have limited our ability to detect small-to-medium effects. In addition, the composition of CAH participants in our study limited our ability to compare salt-wasting vs simple-virilizing forms, as the majority of patients were salt-wasters. Other studies have included mostly patients with the salt-wasting form (15) which could reflect the distribution of salt-wasting to simple-virilizing in clinical CAH populations. Moving forward, it will be imperative for the field to conduct larger studies of patients with CAH, in order to better understand how various clinical features, including commonly seen brain anomalies, may contribute to neurological phenotypes. Previous studies have highlighted an increased prevalence of type 1 Chiari anomalies in patients with CAH (15,39), and treatment regimens as well as adherence to the glucocorticoid and/or mineralocorticoid replacement in CAH varies greatly among patients. Pooled MRI studies across multiple clinical research centers may ultimately help to better understand both individual differences, and common patterns of altered brain structure, by clinical CAH phenotype and concurrent brain anomalies. Lastly, the current study was designed to examine structural differences in *a priori* gray matter in CAH vs control youth. However, studies have also suggested impairments in white matter (15,39) as well as brain function (7,8,10,11,13) in patients with CAH. Thus, studies are currently underway to probe additional brain structural and functional abnormalities as a focus of future research.

We conclude that youth affected with classical CAH have overall smaller intracranial volumes as well as regional atrophy in the prefrontal cortex, amygdala, and hippocampus as compared to healthy controls, suggesting brain alterations are associated with CAH during childhood and adolescent development. Future studies to determine if volumetric differences in the PFC, lateral nucleus of the amygdala, subiculum and CA1 hippocampal subregions map onto physiological and behavioral phenotypes are needed in patients with CAH.

## Acknowledgments

The authors gratefully thank all participants and their families. In addition, we would like to acknowledge Norma Martinez, Heather Ross, Christina Koppin, Kimberly Felix, Veeraya Tanawattanacharoen, and Eva Gabor for assisting with participant recruitment and data collection. Finally, we thank CARES Foundation for their ongoing support of the CHLA CAH Comprehensive Care Center.

## Data Availability Statement

Restrictions apply to the availability of data generated or analyzed during this study to preserve patient confidentiality or because they were used under license. The corresponding author will on request detail the restrictions and any conditions under which access to some data may be provided.

## REFERENCES

1. Merke DP, Bornstein SR. Congenital adrenal hyperplasia. Lancet. 2005;365(9477):2125–2136.

2. Clayton PE, Miller WL, Oberfield SE, Ritzen EM, Sippell WG, Speiser PW. Consensus statement on 21-hydroxylase deficiency from the European Society for Paediatric Endocrinology and the Lawson Wilkins Pediatric Endocrine Society. Horm Res. 2002;58(4):188–195.

3. New MI. Diagnosis and management of congenital adrenal hyperplasia. Annu Rev Med. 1998;49:311–328.

4. DonCarlos LL, Garcia-Ovejero D, Sarkey S, Garcia-Segura LM, Azcoitia I. Androgen receptor immunoreactivity in forebrain axons and dendrites in the rat. Endocrinology. 2003;144(8):3632–3638.

5. Gray JD, Kogan JF, Marrocco J, McEwen BS. Genomic and epigenomic mechanisms of glucocorticoids in the brain. Nat Rev Endocrinol. 2017;13(11):661–673.

6. Mueller SC. Magnetic resonance imaging in paediatric psychoneuroendocrinology: a new frontier for understanding the impact of hormones on emotion and cognition. J Neuroendocrinol. 2013;25(8):762–770.

7. Collaer ML, Hindmarsh PC, Pasterski V, Fane BA, Hines M. Reduced short term memory in congenital adrenal hyperplasia (CAH) and its relationship to spatial and quantitative performance. Psychoneuroendocrinology. 2016;64:164–173.

8. Browne WV, Hindmarsh PC, Pasterski V, Hughes IA, Acerini CL, Spencer D, Neufeld S, Hines M. Working memory performance is reduced in children with congenital adrenal hyperplasia. Horm Behav. 2015;67:83–88.

9. Merke DP, Fields JD, Keil MF, Vaituzis AC, Chrousos GP, Giedd JN. Children with classic congenital adrenal hyperplasia have decreased amygdala volume: potential prenatal and postnatal hormonal effects. J Clin Endocrinol Metab. 2003;88(4):1760–1765.

10. Mazzone L, Mueller SC, Maheu F, VanRyzin C, Merke DP, Ernst M. Emotional memory in early steroid abnormalities: an FMRI study of adolescents with congenital adrenal hyperplasia. Dev Neuropsychol. 2011;36(4):473–492.

11. Ernst M, Maheu FS, Schroth E, Hardin J, Golan LG, Cameron J, Allen R, Holzer S, Nelson E, Pine DS, Merke DP. Amygdala function in adolescents with congenital adrenal hyperplasia: a model for the study of early steroid abnormalities. Neuropsychologia. 2007;45(9):2104–2113.

12. Maheu FS, Merke DP, Schroth EA, Keil MF, Hardin J, Poeth K, Pine DS, Ernst M. Steroid abnormalities and the developing brain: declarative memory for emotionally arousing and neutral material in children with congenital adrenal hyperplasia. Psychoneuroendocrinology. 2008;33(2):238–245.

13. Mueller SC, Daniele T, MacIntyre J, Korelitz K, Carlisi C, Hardin MG, Van Ryzin C, Merke DP, Ernst M. Incentive processing in Congenital Adrenal Hyperplasia (CAH): a reward-based antisaccade study. Psychoneuroendocrinology. 2013;38(5):716–721.

14. Mueller SC, Ng P, Sinaii N, Leschek EW, Green-Golan L, VanRyzin C, Ernst M, Merke DP. Psychiatric characterization of children with genetic causes of hyperandrogenism. Eur J Endocrinol. 2010;163(5):801–810.

15. Webb EA, Elliott L, Carlin D, Wilson M, Hall K, Netherton J, Reed J, Barrett TG, Salwani V, Clayden JD, Arlt W, Krone N, Peet AC, Wood AG. Quantitative Brain MRI in Congenital Adrenal Hyperplasia: In Vivo Assessment of the Cognitive and Structural Impact of Steroid Hormones. J Clin Endocrinol Metab. 2018;103(4):1330–1341.

16. Wan FJ, Swerdlow NR. The basolateral amygdala regulates sensorimotor gating of acoustic startle in the rat. Neuroscience. 1997;76(3):715–724.

17. Schoenbaum G, Chiba AA, Gallagher M. Neural encoding in orbitofrontal cortex and basolateral amygdala during olfactory discrimination learning. J Neurosci. 1999;19(5):1876–1884.

18. Baxter MG, Murray EA. The amygdala and reward. Nat Rev Neurosci. 2002;3(7):563–573.

19. Lacy JW, Yassa MA, Stark SM, Muftuler LT, Stark CE. Distinct pattern separation related transfer functions in human CA3/dentate and CA1 revealed using high-resolution fMRI and variable mnemonic similarity. Learn Mem. 2011;18(1):15–18.

20. Eldridge LL, Engel SA, Zeineh MM, Bookheimer SY, Knowlton BJ. A dissociation of encoding and retrieval processes in the human hippocampus. J Neurosci. 2005;25(13):3280–3286.

21. Adolphs R. Neural systems for recognizing emotion. Curr Opin Neurobiol. 2002;12(2):169–177.

22. Preston AR, Eichenbaum H. Interplay of hippocampus and prefrontal cortex in memory. Curr Biol. 2013;23(17):R764–773.

23. Tyszka JM, Pauli WM. In vivo delineation of subdivisions of the human amygdaloid complex in a high-resolution group template. Human brain mapping. 2016;37(11):3979–3998.

24. Iglesias JE, Augustinack JC, Nguyen K, Player CM, Player A, Wright M, Roy N, Frosch MP, McKee AC, Wald LL, Fischl B, Van Leemput K, Alzheimer’s Disease Neuroimaging I. A computational atlas of the hippocampal formation using ex vivo, ultra-high resolution MRI: Application to adaptive segmentation of in vivo MRI. Neuroimage. 2015;115:117–137.

25. Finkielstain GP, Kim MS, Sinaii N, Nishitani M, Van Ryzin C, Hill SC, Reynolds JC, Hanna RM, Merke DP. Clinical characteristics of a cohort of 244 patients with congenital adrenal hyperplasia. J Clin Endocrinol Metab. 2012;97(12):4429–4438.

26. Kuczmarski RJ, Ogden CL, Guo SS, Grummer-Strawn LM, Flegal KM, Mei Z, Wei R, Curtin LR, Roche AF, Johnson CL. 2000 CDC Growth Charts for the United States: methods and development. Vital Health Stat 11. 2002(246):1–190.

27. Prevention CfDCa. A SAS Program for the 2000 CDC Growth Charts (ages 0 to <20 years). Vol 2019. Atlanta, GA: Division of Nutrition, Physical Activity, and Obesity, National Center for Chronic Disease Prevention and Health Promotion, Centers for Disease Control and Prevention.

28. Greulich WW, Pyle SI. Radiologic Atlas of Skeletal Development of the Hand and Wrist. 2 ed. California: Stanford University Press.

29. Gilsanz V, Ratib O. Hand Bone Age: A Digital Atlas of Skeletal Maturity. Germany: Springer.

30. Wechsler D. Wechsler Adult Intelligence Scale. Fourth ed. San Antonio, TX: Pearson.

31. Reuter M, Schmansky NJ, Rosas HD, Fischl B. Within-subject template estimation for unbiased longitudinal image analysis. Neuroimage. 2012;61(4):1402–1418.

32. Fischl B, Salat DH, Busa E, Albert M, Dieterich M, Haselgrove C, van der Kouwe A, Killiany R, Kennedy D, Klaveness S, Montillo A, Makris N, Rosen B, Dale AM. Whole brain segmentation: automated labeling of neuroanatomical structures in the human brain. Neuron. 2002;33(3):341–355.

33. Backhausen LL, Herting MM, Buse J, Roessner V, Smolka MN, Vetter NC. Quality Control of Structural MRI Images Applied Using FreeSurfer-A Hands-On Workflow to Rate Motion Artifacts. Front Neurosci. 2016;10:558.

34. Avants BB, Duda JT, Zhang H, Gee JC. Multivariate normalization with symmetric diffeomorphisms for multivariate studies. Med Image Comput Comput Assist Interv. 2007;10(Pt 1):359–366.

35. Merke DP, Fields JD, Keil MF, Vaituzis AC, Chrousos GP, Giedd JN. Children with classic congenital adrenal hyperplasia have decreased amygdala volume: potential prenatal and postnatal hormonal effects. J Clin Endocrinol Metab. 2003;88(4):1760–1765.

36. Mnif MF, Kamoun M, Mnif F, Charfi N, Kallel N, Rekik N, Naceur BB, Fourati H, Daoud E, Mnif Z, Sfar MH, Younes-Mhenni S, Sfar MT, Hachicha M, Abid M. Brain magnetic resonance imaging findings in adult patients with congenital adrenal hyperplasia: Increased frequency of white matter impairment and temporal lobe structures dysgenesis. Indian J Endocrinol Metab. 2013;17(1):121–127.

37. Sinforiani E, Livieri C, Mauri M, Bisio P, Sibilla L, Chiesa L, Martelli A. Cognitive and neuroradiological findings in congenital adrenal hyperplasia. Psychoneuroendocrinology. 1994;19(1):55–64.

38. Gaudiano C, Malandrini A, Pollazzon M, Murru S, Mari F, Renieri A, Federico A. Leukoencephalopathy in 21-beta hydroxylase deficiency: report of a family. Brain Dev. 2010;32(5):421–424.

39. Nass R, Heier L, Moshang T, Oberfield S, George A, New MI, Speiser PW. Magnetic resonance imaging in the congenital adrenal hyperplasia population: increased frequency of white-matter abnormalities and temporal lobe atrophy. J Child Neurol. 1997;12(3):181–186.

40. Nunez JL, Huppenbauer CB, McAbee MD, Juraska JM, DonCarlos LL. Androgen receptor expression in the developing male and female rat visual and prefrontal cortex. J Neurobiol. 2003;56(3):293–302.

41. Lupien SJ, McEwen BS. The acute effects of corticosteroids on cognition: integration of animal and human model studies. Brain Res Brain Res Rev. 1997;24(1):1–27.

42. Simerly RB, Chang C, Muramatsu M, Swanson LW. Distribution of androgen and estrogen receptor mRNA-containing cells in the rat brain: an in situ hybridization study. J Comp Neurol. 1990;294(1):76–95.

43. Amaral DG, Price JL, Pikanen A, Carmichael ST. Anatomical organization of the primate amygdaloid complex. In: Aggleton JP, ed. The Amygdala: Neurobiological Aspects of Emotion, Memory, and Mental Dysfunction. New York: Wiley-Liss; 1992:1–66.

44. Duvernoy HM, Francoise C, Risold PY. The Human Hippocampus: Functional Anatomy, Vascularization and Serial Sections with MRI. Springer-Verlag Berlin Heidelberg.

45. H.T. B, M.S. F. Fear and Memory: A View of the Hippocampus Through the Lens of the Amygdala. In: J. DDK, ed. Space, Time and Memory in the Hippocampal Formation. Vienna: Springer; 2014.

46. Raff H, Sharma ST, Nieman LK. Physiological basis for the etiology, diagnosis, and treatment of adrenal disorders: Cushing’s syndrome, adrenal insufficiency, and congenital adrenal hyperplasia. Compr Physiol. 2014;4(2):739–769.

47. Valli I, Crossley NA, Day F, Stone J, Tognin S, Mondelli V, Howes O, Valmaggia L, Pariante C, McGuire P. HPA-axis function and grey matter volume reductions: imaging the diathesis-stress model in individuals at ultra-high risk of psychosis. Transl Psychiatry. 2016;6:e797.

48. O’Mara S. The subiculum: what it does, what it might do, and what neuroanatomy has yet to tell us. J Anat. 2005;207(3):271–282.

49. Tasker JG, Herman JP. Mechanisms of rapid glucocorticoid feedback inhibition of the hypothalamic-pituitary-adrenal axis. Stress. 2011;14(4):398–406.

50. Erickson K, Drevets W, Schulkin J. Glucocorticoid regulation of diverse cognitive functions in normal and pathological emotional states. Neurosci Biobehav Rev. 2003;27(3):233–246.

51. Nebesio TD, Renbarger JL, Nabhan ZM, Ross SE, Slaven JE, Li L, Walvoord EC, Eugster EA. Differential effects of hydrocortisone, prednisone, and dexamethasone on hormonal and pharmacokinetic profiles: a pilot study in children with congenital adrenal hyperplasia. Int J Pediatr Endocrinol. 2016;2016:17.

